# An F-actin mega-cable supports the migration of the sperm nucleus during the fertilization of the polarity-inverted central cell of *Agave inaequidens*

**DOI:** 10.1101/2021.06.23.449664

**Authors:** Alejandra G. González-Gutiérrez, Antonia Gutiérrez-Mora, Jorge Verdín, Benjamín Rodríguez-Garay

## Abstract

Asparagaceae’s large embryo sacs display a central cell nucleus polarized toward the chalaza, which means the sperm nucleus that fuses it during double fertilization migrates a long distance before karyogamy. Because of the size and inverted polarity of the central cell in Asparagaceae, we hypothesize that the second fertilization process is supported by F-actin structures different from the short-range aster-like ones observed in *Arabidopsis*. Here, we analyzed the F-actin dynamics of *Agave inaequidens*, a typical Asparagaceae, before, during, and after central cell fertilization. Several parallel F-actin cables emerging from the nucleus within the central cell, enclosing the vacuole, and reaching the micropylar pole were observed. As fertilization progressed, a thick F-actin mega-cable traversing the vacuole appeared, connecting the central cell nucleus with the micropylar pole near the egg cell. This mega-cable wrapped the sperm nucleus in transit to fuse the central cell one. Once karyogamy finished, the mega-cable disassembled, but new F-actin structures formed during the endosperm development. These observations suggest that Asparagaceae, and probably other plant species with similar embryo sacs, evolved an F-actin machinery specifically adapted to support the migration of the fertilizing sperm nucleus within a large-sized and polarity-inverted central cell.

## Introduction

During angiosperms’ fertilization, two sperm cells are released from the pollen tube at the egg apparatus boundary. One of them fuses the egg cell leading to the first plasmogamy and, subsequently, the first karyogamy that generates the zygote (Hamamura et al., 2011). Afterward, the second sperm fuses with the central cell, and their nuclei lead to the second karyogamy and then further endosperm development (Berger et al., 2008). In *Arabidopsis*, whose central cell nucleus is polarized toward the micropylar end (Sprunck and Gross-Hardt, 2011), the distance the second sperm nucleus travels from the plasmogamy site to the central cell nucleus is around 1 μm (Kawashima and Berger, 2015). However, species in the Asparagaceae family, along with other 13 monocotyledonous families, harbor embryo sacs with a polarity-inverted central cell nucleus, i.e., it localizes near the chalazal pole (Davis, 1966; Bhojwani and Bhatnagar, 1983). In *Agave tequilana,* the distance between the egg cell and the central cell nucleus is about 200-times longer than in *Arabidopsis* (González-Gutiérrez et al., 2014). The latter implies that the second sperm nucleus needs to undertake a longer journey in Asparagaceae. Thus, it is plausible that these plant species evolved a specialized long-range transport machinery to support the migration of the sperm nucleus.

The most accepted model to explain the sperm nuclei transport during fertilization proposes they are carried to the fusion sites by a cytoskeleton supported mechanism (Huang and Russell, 1994; Zhang et al., 1999; Wallwork and Sedgley, 2000; Ye et al., 2002). Kawashima et al. (2014) demonstrated that F-actin, but not microtubules, transports the immotile sperm nuclei during *Arabidopsis* fertilization. It has also been observed the formation of actin structures, called “coronas”, related to double fertilization in *Zea mays* (Huang and Sheridan, 1998) and *Torenia fournieri* (Fu et al., 2000). Similarly, aster-shaped F-actin structures surrounding the sperm nuclei have been observed during *A. thaliana* central cell fertilization (Kawashima et al., 2014).

Based on the evidence described above, we hypothesize F-actin-supported transport of the second sperm nucleus in Asparagaceae may be different from that observed in *Arabidopsis* and other plant models. Since the megagametophyte configuration goes beyond Asparagales, they could also share the same differentiated mechanism. Here, we addressed such hypotheses by characterizing F-actin structures of the *Agave inaequidens* megagametophyte, from the mature embryo sac and sperm nuclear migration at double fertilization to the early endosperm development.

## Results and discussion

### 1. Agave inaequidens harbors a central cell with inverted polarity

To elucidate the mechanism that supports the transport of sperm nuclei during central cell fertilization in Asparagaceae species, we started studying the mature embryo sac of a family member: *A. inaequidens*, so far uncharacterized. *A. inaequidens* displayed the typical mature embryo sac of Asparagales already reported for *A. tequilana* and *A. colimana* (González-Gutiérrez et al., 2014; Barranco-Guzmán et al., 2019). The mature embryo sac (238.58 ±16.28 μm, long; and 128.51±12.20 μm, wide; n=40) was piriform with an haustorial tube at the chalazal end, which contained three antipodal cells (Supp figure 1A and B). Moreover, the embryo sac harbored a large central cell (144.03 ±13.96 μm, long; 124.79±8.89 μm, wide; n=40) with its nucleus located just below the antipodal cells (Supp figure 1A and B). At the opposite side was the egg apparatus (central cell nucleus-egg cell nucleus distance: 156.28±22.62 μm, n=40) composed of an egg (Supp figure 1A and C) and two synergid cells (Supp figure 1A and D). Thus, the female gametophyte development of *A. inaequidens* is of the Polygonum-type (Maheshwari, 1937; 1950) with an atypical central cell nucleus polarity.

### 2. F-actin in the mature embryo sac is restricted to perinuclear and cortical areas

Phalloidin-stained actin filaments were observed attached to the inner membrane of each cell of the mature embryo sac of unpollinated *A. inaequidens* flowers (Figure 1A). The nuclei of these cells were also enveloped by actin filaments from, which some of them extended until reaching the cell periphery (Figure 1A, B, C, and D). Perinuclear actin filaments were scarce in the antipodal cells (Figure 1B) but profuse in the egg apparatus (Figure 1A, C, D). In addition to perinuclear actin, synergids displayed a pronounced aggregation of actin filaments at the micropylar end, around the space occupied by their nuclei (Figure 1C). A similar F-actin arrangement, but oriented to the chalazal pole, was observed around the egg cell nucleus (Figure 1D). Actin and other cytoskeleton proteins provide the support to polarize and anchor nuclei within a cell, which is developmentally programed and associated with the final cellular function (Starr and Han, 2003; Smith, 2003; Gu et al., 2005).

**Figure 1.**
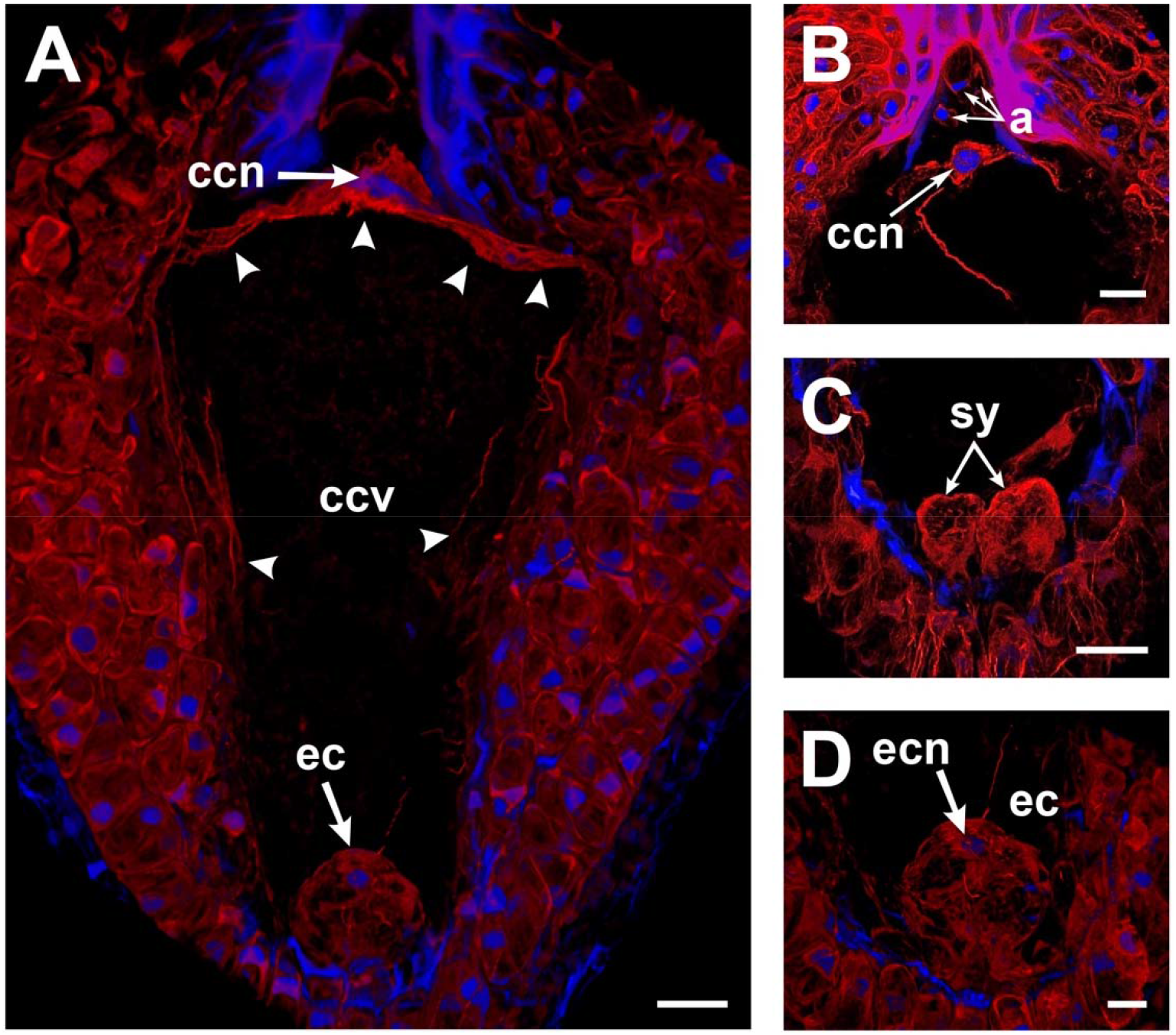
Perinuclear and cortical F-actin in *Agave inaequidens* mature embryo sac. **(A)** Rhodamine-phalloidin-stained actin filaments are located at the periphery of the central cell. **(B)** The central cell nucleus displays a dense perinuclear F-actin coat, while it is scarce in the antipodal cells. **(C)** F-actin is denser at the synergids’ micropylar end, where their nuclei are located. **(D)** Egg cell cortical and perinuclear actin filaments. ccn=central cell nucleus, ccv=central cell vacuole, a=antipodal cells, sy=synergids, ec=egg cell, ecn=egg cell nucleus. Arrow heads=cortical actin filaments of the central cell. All micrographs are oriented with the chalazal pole at the top. Bar in (A-C) =20 μm and (D) =10 μm.

Most of the space in the central cell was occupied by a large vacuole (Figure 1A), while other cytoplasmic content, including cortical actin filaments, was restricted to the cell periphery (Figure 1A). Studies in *Arabidopsis* root epidermal and egg cells and Tobacco somatic BY-2 cells showed that the size and dynamics of vacuoles are F-actin-dependent (Sheuring et al., 2016; Higaki et al., 2006; Kimata et al., 2016). Moreover, it has been suggested that the cell size in plants is regulated via the vacuole size, which prevents a drastic modification of the cytoplasmic content (Sheuring et al., 2016).

In addition to cortical F-actin, actin filaments extending from the central cell nucleus seemed to attach it to the chalazal area (Figure 1A). The nuclear positioning of the central cell by actin filaments was demonstrated by Kawashima and Berger (2015), who disrupted central cell F-actin in *A. thaliana* mature embryo sacs causing a shift of the central cell nucleus from its micropylar to a central position.

### 3. F-actin cables projected from the central cell nucleus form a tunnel-like structure before fertilization

To observe changes in actin cytoskeleton during double fertilization in *A. inaequidens*, flowers were hand-pollinated and collected at different hours after pollination (HAP). In female gametophytes processed at 24 and 30 HAP, actin filaments at the central cell micropylar end started to project from the cortical area to the cell center, and an arch-shaped accumulation of filaments could be seen close to the egg apparatus (Figure 2A). Simultaneously, the F-actin coat of the central cell nucleus and the cables that anchored it to the inner membrane began to extend towards the middle part of the cell, forming thick F-actin cables parallel to the chalazal-micropillar axis (Figure 2B). Finally, around 32-36 HAP, those F-actin cables reached the micropylar end, in the vicinity of the egg apparatus, building a structure we named “actin tunnel” (Figure 2C). This actin tunnel seems to be the functional equivalent of the actin cables that extend from the micropylar to the chalazal region of *Arabidopsis* central cell, which shows an inward movement associated with the sperm nuclear migration (Kawashima et al., 2015).

**Figure 2.**
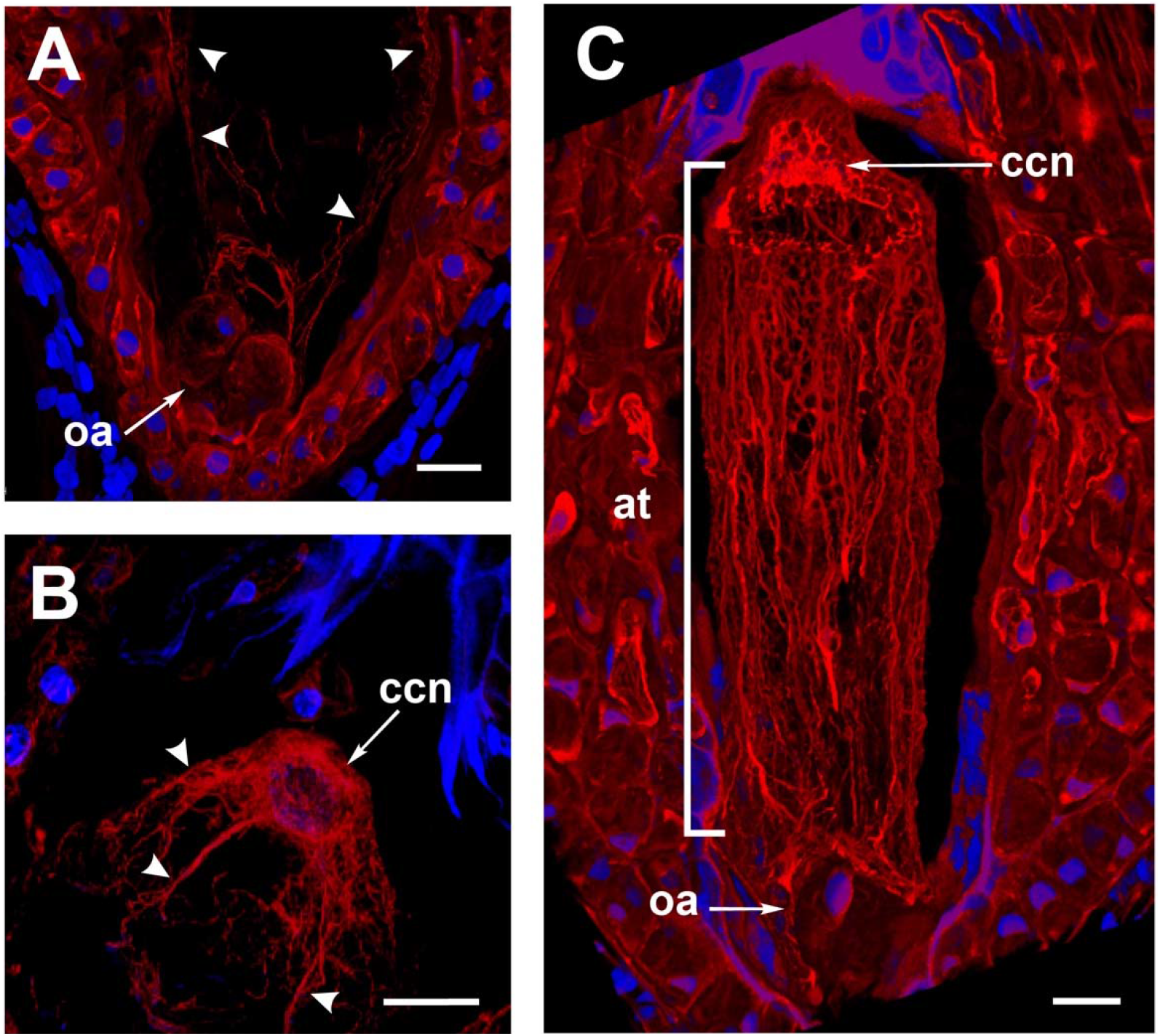
The formation of the *Agave inaequidens* actin-tunnel. **(A)** Central cell F-actin starts nucleating at the cell body. **(B)** Actin filaments projected from the central cell nucleus towards the embryo sac micropylar pole. **(C)** Actin-tunnel formed by several parallel filaments extends from the central cell nucleus to the micropylar pole, next to the egg cell. oa=ovular apparatus, ccn=central cell nucleus, at=actin tunnel. Arrowheads in (A) and (B) = actin filaments. In all cases, F-actin was stained with rhodamine-phalloidin. Nuclei were stained with Hoechst 33258. All micrographs are oriented with the chalazal pole at the top. Bar in (A-C) =20 μm.

The deployment of these F-actin networks was evident only in ovules of pollinated flowers whose pollen tubes were close to the micropyle. Therefore, the changes shown by the actin cytoskeleton in the embryo sac correlate with the release of the two sperm cells into the embryo sac (Weterings and Russell, 2004).

Even though it is clearly associated with the fertilization process, the exact physiological role of the actin tunnel is intriguing. Because the vacuolar dynamics have been demonstrated to be mediated by actin filaments, it is plausible that the actin tunnel is involved in remodeling the central cell vacuole.

### 4. An F-actin mega-cable supports the sperm nucleus migration during the central cell fertilization

Feulgen-stained ovules in which the pollen tube had already arrived at one of the synergids displayed a “central strand” traversing the central cell vacuole (Figure 3A). This central strand was putatively composed of cytoplasm (trans-vacuolar strand) and connected the central cell nucleus directly to the micropylar end at the egg cell boundary (Figure 3A).

**Figure 3.**
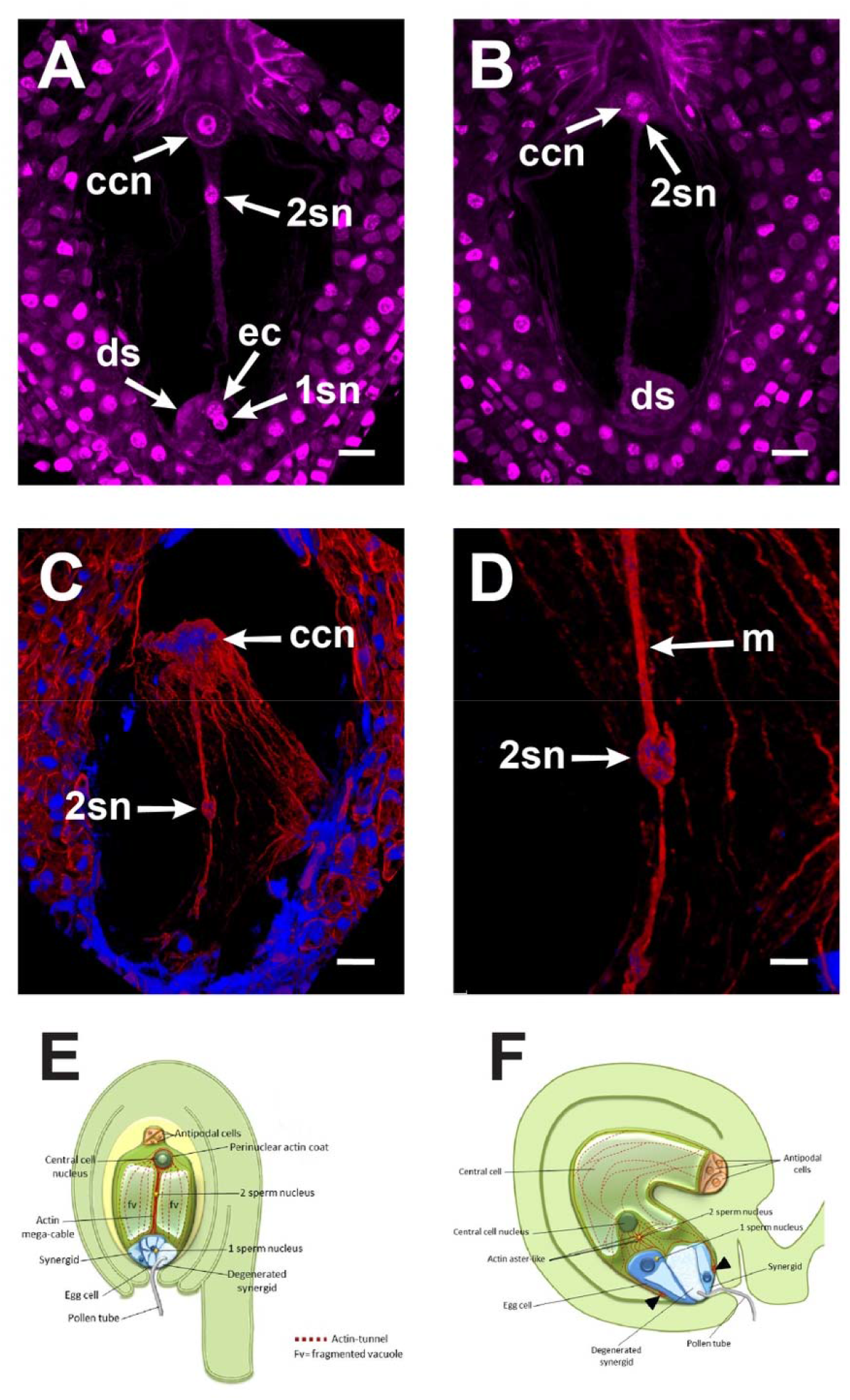
Central cell fertilization of *Agave inaequidens*. **(A)** A cytoplasmic transvacuolar strand and the second sperm nucleus revealed by Feulgen staining were observed at the central cell. **(B)** Second sperm nucleus getting close for plasmogamy/karyogamy. **(C and D)** The rhodamine-phalloidin stained actin mega-cable traversing the central cell vacuole wraps the second sperm nucleus (stained with Hoechst 33258). **(E)** Model of the central cell fertilization in *Agave* embryo sac where an actin-based mega-cable traverses the central vacuole, wraps the sperm nucleus, and supports its migration for the second karyogamy event. **(F)** In *Arabidopsis*, during the second fertilization, the sperm nucleus is surrounded by an aster-shaped structure that moves it toward the central cell one (Modified from Dresselhaus et al., 2016). ccn=central cell nucleus, 2sn=second sperm nucleus, ec=egg cell, ds=degenerated synergid, m=actin mega-cable. All micrographs are oriented with the chalazal pole at the top. Bar in (A-C) =20 μm and (D) =10 μm.

One of the sperms performed plasmogamy and its nucleus fused with the egg cell nucleus. Meanwhile, the second sperm cell and the central cell fused their membranes. Then, the sperm nucleus started a journey through the large central cell vacuole moving along the central trans-vacuolar strand to get the central cell nucleus at the opposite side of the embryo sac (Figure 3A, B). In *Torenia fournieri,* a thick cytoplasmic strand appeared above the ovular apparatus approximately 15 HAP and five hours after karyogamy of the first sperm nucleus with the egg cell one (Higashiyama et al., 1997). Transvacuolar strands also aid in the mobility of polar nuclei, which fuse to form the central cell nucleus (Susaki et al., 2021).

Just before the second fertilization took place, a new thick F-actin cable, that we named “mega-cable”, extended from the actin coat of the central cell nucleus to the micropylar pole of the cell, more precisely at the boundary of the central cell with the egg cell (Figure 3C, D). It is unclear whether the mega-cable evolved from one or several of the pre-existing tunnel-forming cables or emerged *de novo* as a specialized structure.

The mega-cable encompassed the sperm nucleus in transit to the central cell nucleus (Figure 3C, D). Because of the similar position of the cytoplasmic strand and the F-actin mega-cable within the embryo sac, it is reasonable to hypothesize that the latter fills the space created by the cytoplasmic strand to allow the transit of the sperm nucleus. It is well known that actin filaments are involved in cytoplasmic streaming and that cytoplasmic strands function as transport routes for proteins and organelles (Shimmen and Yokota, 2004). In *A. inaequidens*, the actin mega-cable seems to be the functional equivalent of the actin track, and the aster-like structure generated in *Arabidopsis* associated with the migration of the second sperm nucleus during central cell fertilization (Kawashima et al., 2014). Nevertheless, due to the differences between both structures, the mechanistic implications are also different. While the actin track and the aster-like structure are pleomorphic and do not connect the sites of plasmogamy and karyogamy, the mega-cable establishes a continuous connection between the central cell nucleus and its micropylar end, where most probably the second plasmogamy occurs (Figure 3C). There, the sperm nucleus may be taken and actively transported by the mega-cable until it gets in touch with the central cell nucleus (Figure 3D). The comparative diagrams summarize the principal differences and similarities between F-actin mechanisms at the second fertilization of *Agave* (Figure 3E) and *Arabidopsis* (Figure 3F). Our observations demonstrate a direct association of the actin mega-cable with the second sperm nucleus migration; nevertheless, the precise mechanism by which the nucleus is transported remains to be determined.

The actin tunnel and the mega-cable were disassembled once the migration of the second sperm nucleus and karyogamy with the central cell nucleus occurred. The latter implies that these structures are mainly formed to participate in the fertilization process. Afterwards, the primary endosperm nucleus was formed, and, along with it, new F-actin structures were built (Supp figure 2A and B) (González-Gutiérrez, to be published elsewhere).

Here we showed that specialized F-actin structures, actin tunnel and mega-cable, are clearly related to *A. inaequidens* sperm nuclei migration for the central cell fertilization. *inaequidens*, and Asparagaceae in general, are characterized by particularly large central cells whose nuclei, in addition, are polarized to the chalaza. Our observations suggest the actin tunnel, but especially the actin-mega cable, might be an evolutionary solution in these plant species to the challenge of transporting an immotile sperm nucleus a long distance. Despite the actin tunnel and the mega cable seem to have functional analogs in *Arabidopsis* central cell fertilization (the track and the aster-like structure), their structure and functional scope are clearly different. Why is the fertilizing sperm nucleus in *A. inaequidens* not moved by an aster-like structure? We hypothesize that the differences in F-actin structures developed on each system depend on the distance that the sperm nucleus needs to be transported. Actin structures adopt different configurations depending on the distance the cargo needs to be transported in plant cells. Individual or thin actin filaments are associated with short-range cargo targeting, while thicker actin cables are necessary for long-distance transport (Geitmann and Emons, 2000). Because of its thin-cable configuration, an aster-like structure might be more convenient for a short-range movement Thus, as observed, a robust mega-cable seems a better solution for a long journey.

Although our staining methodology did not allow either higher-resolution or live imaging, our observations suggest that the fertilizing sperm nucleus is wrapped by the F-actin mega-cable, implying that the sperm nucleus is not moved by an actin-associated motor protein. Instead, the sperm nucleus might be transported together with the mega-cable by a treadmilling mechanism. Nevertheless, the sperm nucleus transporting mechanism should be further investigated.

Our observations describe for the first time the adaptation of a critical cellular process, central cell fertilization, to the particularities of a minority but important group of plant species, so far only accessible by staining techniques since most of them remain genetically difficult to manage.

## Methods

### Plant material

Inflorescences with mature flowers from 20 plants of *A. inaequidens* were collected in Atemajac de Brizuela, Jalisco, Mexico, during the flowering seasons (May-June) of 2017-2020.

### Pollination and collection of specimens

Inflorescences were maintained at the laboratory in freshwater. Flowers were emasculated at anthesis and covered with glassine paper bags to avoid free pollination. Extracted anthers were kept in Petri dishes at 4°C until dehiscence. At this time, mature pollen grains were recovered from the anthers and tested for viability using the *in vitro* method for pollen germination of the *Agave* genus, proposed by López-Díaz and Rodríguez-Garay (2008).

Once the stigmas of emasculated flowers were receptive (presence of a pollination drop), ten flowers per inflorescence were selected and processed as described below. Flowers collected at this stage were considered “time 0”. The remaining flowers at this developmental stage were hand-pollinated (cross-pollination) and collected at different times between 1 and 72 hours after pollination (HAP), so that it was possible to record actin cytoskeleton dynamics in mature *A. inaequidens* embryo sacs during double fertilization and the first endosperm divisions. Ovules of the same flower were dissected with fine-point tweezers and an insulin needle under the stereoscope and evenly divided into two centrifuge tubes to be processed with the histological techniques described below.

Only “normal” megagametophytes were taken into account, i.e., piriform embryo sacs with a pronounced haustorial tube and the four cellular types contained in seven cells. Collapsed embryo sacs and embryo sacs lacking any cellular type due to abnormal growth of the nucellar tissue were discarded. At least 500 ovules encompassing the different developmental stages were analyzed with each staining technique.

### Feulgen staining (Barrell and Grossniklaus, 2005)

Although Feulgen staining primarily binds to DNA (Kalinowska and Dresselhaus, 2020), some other structures like cell walls (Barrell and Grossniklaus, 2005) and cytoplasm could be weakly stained (Chieco and Derenzini, 1999).

Feulgen staining adapted with minor modifications for agave ovules was used as a massive method for analyzing the general development stage of the embryo sacs. In short, after fixation in FAA (10:5:50:35 formaldehyde: acetic acid: ethanol: distilled water) for 24 h and kept overnight in 70% ethanol, 4°C, ovules were treated with 1 M HCl for 1:30 h, 5.8 M HCl for 2 h and, 1 M HCl for 1 h at room temperature. Subsequently, ovules were rinsed three times with distilled water and stained with Schiff reagent (Sigma cat. no. S5133) for 3 h at room temperature. Dehydration was carried out by an increasing concentration series of 30, 50, 70, 90, 95, 100% ethanol for 30 min, and an additional 100% ethanol incubation for 30 min. Finally, ovules were clarified by a series of methyl salicylate:ethanol solutions of 3:1, 1:1, 1:3, for one hour. For observation, samples were mounted in 100% methyl salicylate and examined on a Leica TCS SPE confocal microscope at Ex=532 nm and Em=555-700 nm. Images were acquired and processed with LAS X® software (Leica Microsystems).

### F-actin whole-mount staining (González-Gutiérrez et al., 2020)

Ovules previously dissected were pre-incubated in ASB (Actin Stabilizing Buffer) (50 mM PIPES, 10 mM EGTA, and 1 mM MgCl2, pH 6.8 adjusted with 10M KOH) at 55° C for 5 minutes. Then, ovules were fixed with 4% formaldehyde in ASB for 10 min at room temperature (25° C). Afterward, ovules were washed twice with ASB. Two quick rinses with acetone (−20° C), followed by a 5 min incubation in acetone (−20° C), were performed for cuticle solubilization and membrane permeabilization. After this time elapsed, acetone was removed from the microtubes, and ovules were washed 3 times with ASB until the solution remained crystalline. Then, ovules were incubated in blocking solution (1% BSA in ASB) for 20 min at room temperature and stained with 0.33 mM rhodamine-phalloidin and 3 mg/ml Hoechst 33258 (diluted in blocking solution), overnight at 4°C. Before clarification, ovules were dehydrated in an increasing concentration series of isopropanol (diluted in ASB) at 4°C, for 7 min each: 75, 85, 95,100%, and an additional 100% isopropanol step for 12 min. Tissue clarification was carried out by adding 1:1 methyl salicylate-isopropanol solution until all ovules were precipitated at the microtube bottom. Finally, ovules were incubated in 100% methyl salicylate for at least 30 min before observation. Samples were analyzed under a Leica TCS SPE confocal microscope using a 532 nm laser for rhodamine-phalloidin (ex/em = 540/556 nm) and a 405 nm laser for Hoechst 33258 observation (ex/em = 352/461 nm). Images were taken and managed with the LAS X® software (Leica Microsystems).

## Acknowledgments

We thank Rosa Isela Martínez-Contreras for technical support, José Aldana-Padilla for diagrams draw, and Hiram Rodríguez-Julián for figures layout arrangement.

---

**Supplementary figure 1.**
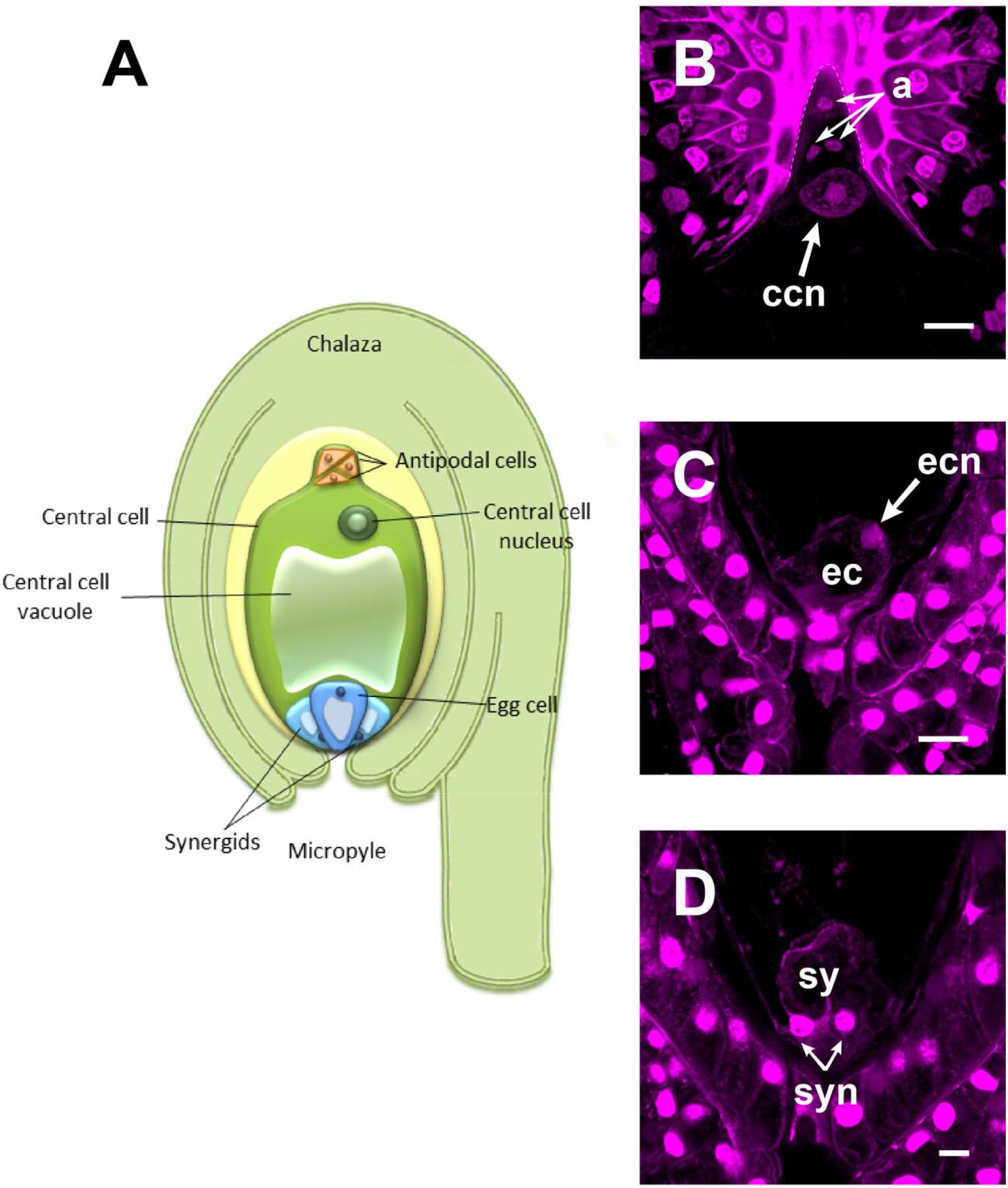
The mature female gametophyte of *Agave inaequidens*. **(A)** Scheme of a seven-celled embryo sac arranged in four cell types (Feulgen stained): three antipodal cells, placed in the haustorial tube, and a large central cell with its nucleus polarized toward the chalaza. **(B)** At the micropylar end the egg apparatus is composed of an egg cell **(C)** and two synergids **(D).** a=antipodal cells, ccn=central cell nucleus, ec=egg cell, ecn=egg cell nucleus, sy=synergids, syn=synergids nuclei. Dashed line in (B) shows the haustorial tube. Barr in (B-C) = 20 μm and (D) =10 μm.

**Supplementary figure 2.**
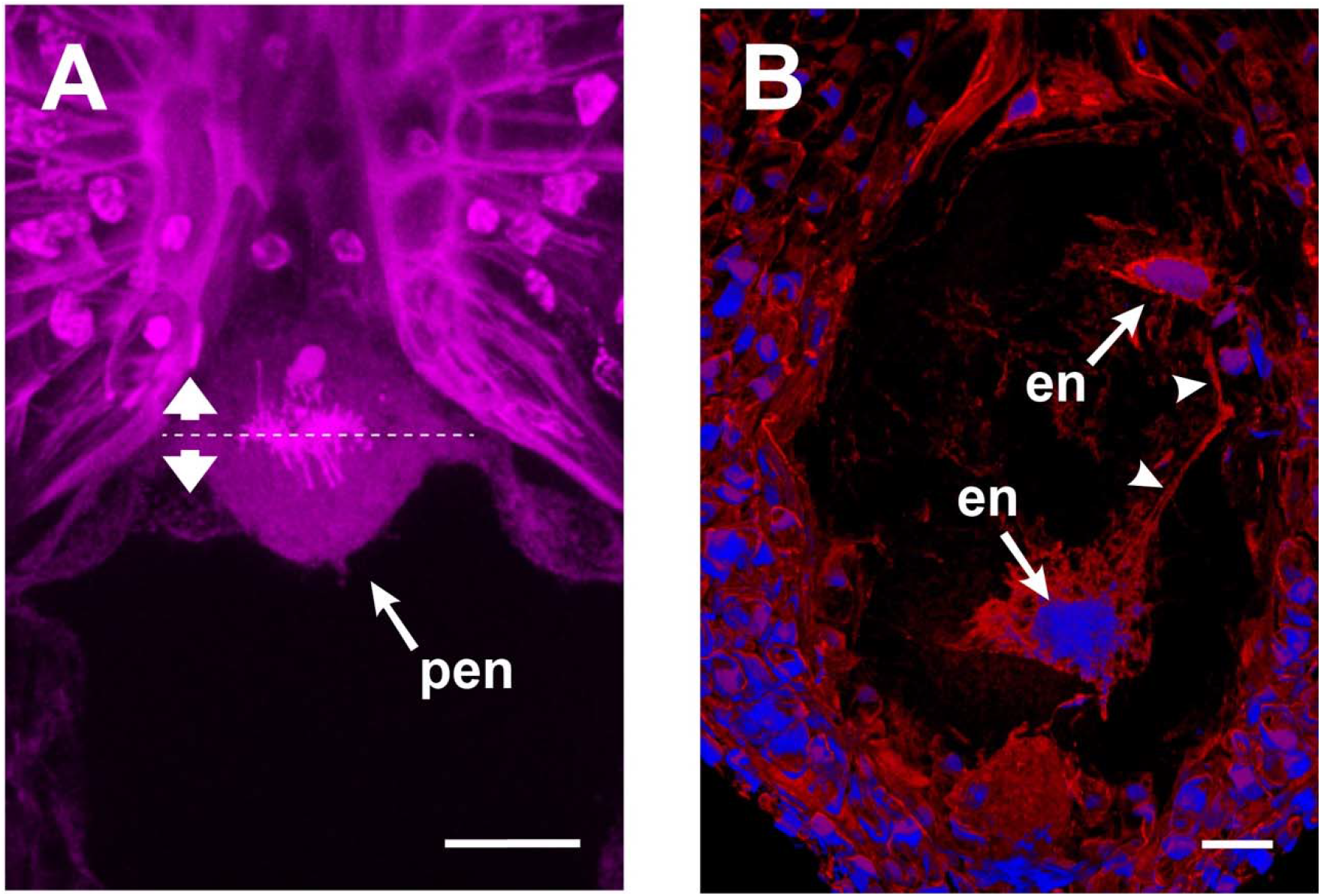
F-actin structures in the developing endosperm of *Agave inaequidens*. **(A)** Dividing primary endosperm nucleus stained with Feulgen. **(B)** F-actin filaments (arrowheads) connecting endosperm nuclei in the coenocyte. F-actin was stained with rhodamine-phalloidin; nuclei were stained with Hoechst 33258. pen=primary endosperm nucleus, en=endosperm nuclei. The dashed line in (A) indicates the division plane of the pen to form the first two daughter nuclei. Micrographs are oriented with the chalazal pole at the top. Bar in (A) and (B) =20 μm.

## References

Barranco-Guzmán, A. M., González-Gutiérrez, A. G. and Rodríguez-Garay B. (2019). The embryo sac development of *Manfreda elongata* (Asparagaceae). Flora 260, 151480. https://doi.org/10.1016/j.flora.2019.151480

Barrell, P. J. and Grossniklaus, U. (2005). Confocal microscopy of whole ovules for analysis of reproductive development: the *elongate1* mutant affects meiosis II. Plant J., 43, 309–320. doi: 10.1111/j.1365-313X.2005.02456.x.

Berger, F., Hamamura Y., Ingouff M. and Higashiyama T. (2008). Double fertilization caught in the act. Trends Plant Sci. 13, 437–443. doi: 10.1016/j.tplants.2008.05.011.

Bhojwani, S.S. and Bhatnagar, S. P. (1983). The embryology of angiosperms. Vikas Publishing House Pvt. Ltd., New Delhi, India.

Chieco, P. and Derenzini, M. (1999). The Feulgen reaction 75 years on. Histochem. Cell Biol. 111, 345–358. https://doi.org/10.1007/s004180050367.

Davis, G. L. (1966). Systematic Embryology of the Angiosperms. Wiley, New York.

Dresselhaus, T., Sprunck, S. and Wessel, G. M. (2016). Fertilization mechanisms in flowering plants. Curr Biol. 26, R125–R139. doi: 10.1016/j.cub.2015.12.032.

Fu, Y., Yuan, M., Huang, B. Q., Yang, H. Y., Zee, S. Y. and O’Brien, T. P. (2000). Changes in actin organization in the living egg apparatus of *Torenia fournieri* during fertilization. Sex. Plant Reprod. 12, 315–322. https://doi.org/10.1007/s004970000026

Geitmann, A. and Emons, A. M. C. (2000). The cytoskeleton in plant and fungal cell tip growth. J Microsc. 198, 218–245. doi: 10.1046/j.1365-2818.2000.00702.x.

González-Gutiérrez, A. G., Gutiérrez-Mora, A. and Rodríguez-Garay, B. (2014). Embryo sac formation and early embryo development in *Agave tequilana* (Asparagaceae). SpringerPlus 3, 575. doi: 10.1186/2193-1801-3-575.

González-Gutiérrez, A. G., Verdín, J. and Rodríguez-Garay, B. (2020). Simple whole-mount staining protocol of F-Actin for studies of the female gametophyte in Agavoideae and other crassinucellate ovules. Front. Plant Sci. 11, 384. https://doi.org/10.3389/fpls.2020.00384.

González-Gutiérrez, A.G. and Rodríguez-Garay, B. (2016). Embryogenesis in *Polianthes tuberosa* L var. Simple: from megasporogenesis to early embryo development. SpringerPlus 5,1804. https://doi.org/10.1186/s40064-016-3528-z.

Gu, Y., Fu, Y., Dowd, P., Li, S., Vernoud, V., Gilroy, S. and Yang, Z. (2005). A Rho family GTPase controls actin dynamics and tip growth via two counteracting downstream pathways in pollen tubes. J. Cell Biol. 169, 127–138. doi: 10.1083/jcb.200409140.

Hamamura, Y., Saito, C., Awai, C., Kurihara, D., Miyawak, iA., Nakagawa, T., Kanaoka, M. M., Sasaki, N., Nakano, A., Berger, F. and Higashiyama, T. (2011). Live-cell imaging reveals the dynamics of two sperm cells during double fertilization in *Arabidopsis thaliana*. Curr. Biol. 21, 497–502. doi: 10.1016/j.cub.2011.02.013.

Higaki, T., Kutsuna, N., Okubo, E., Sano, T. and Hasezawa, S. (2006). Actin microfilaments regulate vacuolar structures and dynamics: dual observation of actin microfilaments and vacuolar membrane in living tobacco BY-2 cells. Plant Cell Physiol. 47, 839–852. doi: 10.1093/pcp/pcj056.

Huang, B. Q. and Sheridan, W. F. (1998). Action coronas in normal and indeterminate gametophyte1 embryo sacs of maize. Sex Plant Reprod. 11, 257–264. doi: 10.1105/tpc.8.8.1391.

Huang, B. Q. and Russell S. D. (1994). Fertilization in *Nicotiana tabacum*: Cytoskeletal modifications in the embryo sac during synergid degeneration. A hypothesis for short-distance transport of sperm cells prior to gamete fusion. Planta 195, 200–214. https://doi.org/10.1007/BF01101679

Kalinowska K., Chen J. and Dresselhaus T. (2020) Imaging of embryo sac and early seed development in Maize after Feulgen staining. In: Bayer M. (eds) Plant Embryogenesis. Methods in Molecular Biology, vol 2122. Humana, New York, NY. https://doi.org/10.1007/978-1-0716-0342-0_14.

Kawashima, T., Maruyama, D., Shagirov, M., Li, J., Hamamura, Y., Yelagandula, R. and Berger, F. (2014). Dynamic F-actin movement is essential for fertilization in *Arabidopsis thaliana*. eLife 3, e04501. doi: 10.7554/eLife.04501.

Kawashima, T and Berger, F. (2015). The central cell nuclear position at the micropyle is maintained by the balance of F-actin dynamics, but dispensable for karyogamy in Arabidopsis. Plant Reprod. 28,103–10. doi: 10.1007/s00497-015-0259-1.

Kimata, Y., Higaki, T., Kawashima, T., Kurihara, D., Sato, Y., Yamada, T. and Ueda, M. (2016). Cytoskeleton dynamics control the first asymmetric cell division in *Arabidopsis* zygote. P. Natl. Acad. Sci. USA, 113, 14157–14162. https://doi.org/10.1073/pnas.1613979113.

López-Díaz, S. and Rodríguez-Garay B. (2008). Simple methods for *in vitro* pollen germination and pollen preservation of selected species of the genus *Agave*. e-Gnosis 6, 1–7.

Maheshwari, P. (1937). A critical review of the types of embryo sacs in angiosperms. New Phytol. 36, 359–417.

Maheshwari, P. (1950). An introduction to the embryology of angiosperms. McGraw-Hill, New York, pp: 65–67.

Scheuring, D., Löfke, C., Krüger, F., Kittelmann, M., Eisa, A., Hughes, L., Smith, R. S., Hawes, C., Shumacher, K. and Kleine-Vehn, J. (2016). Actin-dependent vacuolar occupancy of the cell determines auxin-induced growth repression. PNAS USA 113, 452–457. https://doi.org/10.1073/pnas.1517445113.

Shimmen, T. and Yokota, E. (2004). Cytoplasmic streaming in plants. Curr. Opin. Cell Biol. 16, 68–72. https://doi.org/10.1016/j.ceb.2003.11.009.

Smith, L. G. (2003). Cytoskeletal control of plant cell shape: getting the fine points. Curr. Opin. Plant Biol. 6, 63–73. doi: 10.1016/s1369-5266(02)00012-2.

Sprunck, S. and Gross-Hardt, R. (2011). Nuclear behavior, cell polarity, and cell specification in the female gametophyte. Sex Plant Reprod. 24, 123–136. doi: 10.1007/s00497-011-0161-4.

Starr, D. A. and Han, M. (2003). ANChors away: an actin based mechanism of nuclear positioning. J. Cell Sci. 116, 211–216. https://doi.org/10.1242/jcs.00248.

Susak,i D., Suzuki, T., Maruyama, D., Ueda, M., Higashiyama, T. and Kurihara, D. (2021). Dynamics of the cell fate specifications during female gametophyte development in *Arabidopsis*. PLoS Biol. 19, e3001123. https://doi.org/10.1371/journal.pbio.3001123.

Wallwork, M. A. B. and Sedgley, M. (2000). Early events in the penetration of the embryo sac in *Torenia fournieri* (Lind.). Ann. Bot. 85, 447–454. https://doi.org/10.1006/anbo.1999.1093.

Wetering, K. and Russell, S. D. (2004). Experimental analysis of the fertilization process. Plant Cell 16, S107–S118. https://doi.org/10.1105/tpc.016873.

Ye, X.-L., Yeung, E. C. and Zee, S.-Y. (2002). Sperm movement during double fertilization of a flowering plant, *Phaius tankervilliae*. Planta 215, 60–66. doi: 10.1007/s00425-002-0736-2.

